# Impact of chromosome fusions on 3D genome organization and gene expression in budding yeast

**DOI:** 10.1101/237263

**Authors:** Marco Di Stefano, Francesca Di Giovanni, Davide Baù, Lucas B. Carey, Marc A. Marti-Renom, Manuel Mendoza

## Abstract

The three-dimensional organization of chromosomes can influence transcription. However, the frequency and magnitude of these effects remains debated. To determine how changes in chromosome positioning affect transcription across thousands of genes with minimal perturbation, we characterized nuclear organization and global gene expression in budding yeast containing chromosome fusions. We used computational modelling and single cell imaging to determine chromosome position and integrated these data with genome-wide transcriptional profiles from RNA sequencing. We find that chromosome fusions dramatically alter 3D nuclear organization without leading to strong genome-wide changes in transcription. However, we observe a mild but significant and reproducible increase in expression of genes near fusion sites. Modeling suggests that this is due to both disruption of telomere-associated silencing and the displacement of genes relative to the nuclear periphery. A 10% decrease in the predicted time a gene spends near the nuclear periphery is associated with a 10% increase in expression. These data suggest that basal transcriptional activity is sensitive to radial changes on gene position, and provide insight into the functional relevance of budding yeast chromosome-level three-dimensional organization in gene expression.

## INTRODUCTION

Chromosomes in interphase nuclei are spatially distributed in a non-random manner. Indeed, chromosomes are organized in distinct structural units and their organization influences nuclear functions such as transcription, replication and DNA damage repair (reviewed in [1-4]). In animal cells individual chromosomes tend to occupy defined nuclear regions termed “chromosome territories” (CT) [5-8]. In animal cells, the spatial distribution of CTs can be size- and gene density-dependent; in several cell types, gene-poor chromosomes associate preferentially with the nuclear periphery, whereas gene-rich chromosomes are enriched in the nuclear interior [9,10]. In addition, distinct structural domains at the sub-chromosomal level have been identified by microscopy, termed chromosomal domains, or CDs [11]. Chromosomal domains may correspond to sub-chromosomal units defined by their increased interaction frequencies with each other or with the nuclear lamina. Notably, individual genes can display mobility within chromosomal and sub-chromosomal domains, and this has been correlated with changes in their expression level during cell differentiation [12]. It remains unclear, however, if the position of individual genes within the nucleus affects their expression and/or their ability to be silenced or activated in response to different stimuli, or if these expression-related properties are merely correlated with spatial organization.

Studies in the budding yeast *Saccharomyces cerevisiae* have provided insight into the functional role of nuclear spatial organization (reviewed in [13-15]). In this organism, chromosome organization is highly stereotypical. The 16 centromeres localize around the spindle pole body (SPB, the equivalent of the animal cell centrosome); whereas the 32 telomeres cluster in 3-8 different foci at the nuclear periphery. Chromosome arms thus extend away from the SPB towards the nuclear periphery where telomeres are anchored, and their specific distribution is linked to their length. Finally, the nucleolus is positioned on the opposite side of the SPB, and is organized around 100-200 repeats of ribosomal DNA located in chromosome XII. Certain aspects of nuclear organization can have an impact in gene expression in budding yeast. On one hand, artificial tethering of reporter genes to subtelomeric regions and to the nuclear periphery can lead to their silencing [16-19]. Moreover, perinuclear tethering of the *CLN2* cyclin gene in daughter cells mediates its silencing during the G1 phase [20]. The association of silent information regulators (SIR) factors with telomeres also contributes to perinuclear repression [19]. Accordingly, genes within 20 kb from telomeres are poorly expressed, and this depends at least partially on SIR proteins and telomere anchoring to the nuclear periphery [19,21]. On the other hand, some inducible genes translocate from the nuclear interior to the periphery upon activation, where they interact with nuclear pore complexes [22-26], and artificial targeting of genes to nuclear pores can also lead to their transcriptional activation [26-28]. Thus, the yeast nuclear periphery appears to harbor transcriptionally repressing and activating domains. How the three-dimensional organization of the yeast genome shapes global transcription levels remains largely unexplored.

To study the effect of nuclear organization on transcription in budding yeast, we took advantage of previously described strains bearing Fusion Chromosomes (FCs) [29,30]. Here, we show that FC strains have a grossly altered nuclear organization in interphase that is not associated with dramatic genome-wide changes in transcription. However, displacement of fusion chromosome genes away from the nuclear periphery does lead to mild, but consistent and reproducible changes in expression across a large number of genes; on average a 10% shift away from the nuclear periphery leads to a 10% increase in expression. These effects are associated with both disruption of telomere-associated silencing and with displacement away from the nuclear periphery. These results suggest that radial chromosome-level spatial organization plays a role in transcriptional regulation in budding yeast. Furthermore, this study demonstrates that FC strains are an excellent experimental system in which to test the functional relevance of nuclear organization, and the global role of chromosomal rearrangements on various aspects of cell physiology, such as DNA replication timing and DNA damage repair.

## RESULTS

### A computational model to study the impact of yeast nuclear organization in gene expression

To study how the three-dimensional organization of the genome affects gene expression, we first sought to establish how gene position correlates with transcription levels in wild-type budding yeast cells. To estimate gene position we built computational models of chromosomes in the interphase G1 nucleus, a strategy that has proven useful in recapitulating chromosome-level nuclear organization in budding yeast [31-33]. We modelled chromosomes as bead-and-spring chains, an approach previously validated for modelling the general physical properties of chromatin fibers [34,35]. Details on the polymer modelling are found in *Materials and Methods* and summarized in Figure 1A. Briefly, chromosomes were confined inside a sphere of 2 μm in diameter corresponding to the interphase nuclear size. Centromeres were confined in a spherical region of radius 150 nm at one pole of the nuclear sphere to account for the tethering of centromeres to the spindle pole body by microtubules [36]. The dynamic association of telomeres with the nuclear envelope was modelled with the periphery of the sphere attracting the terminal beads of chromosome chains. Finally, to reproduce the confinement of the rDNA in the nucleolus, the particles corresponding to rDNA were restrained to a region located at the opposite side of the SPB. An ensemble of chromosomal polymer models was generated using Brownian motion dynamics. A total of 10,000 model conformations satisfying all the imposed restraints were then selected and analyzed for the likelihood of particular loci and chromosomes to be positioned in specific regions of the cell nucleus (Figure 1B).

**Figure 1.**
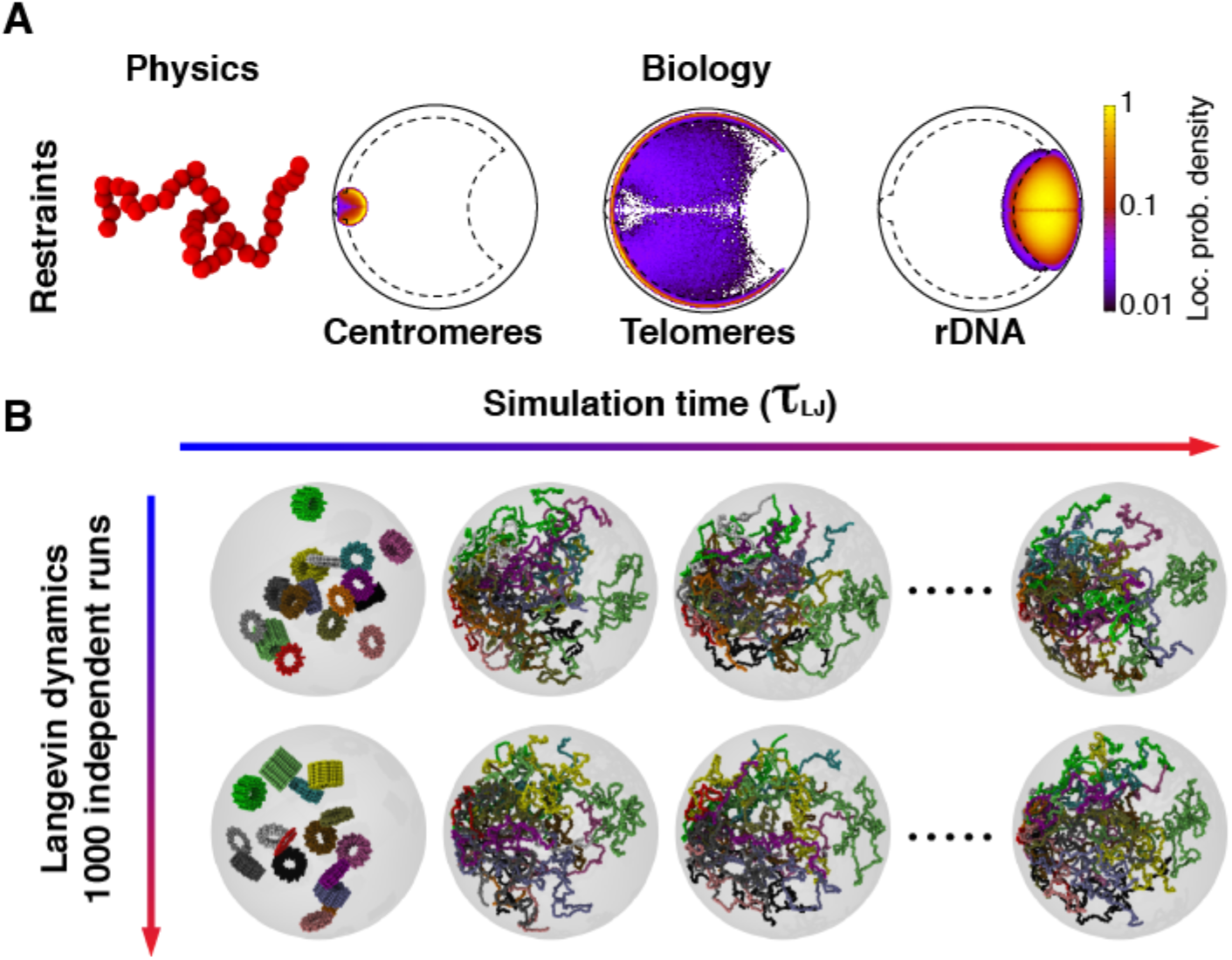
Computational modelling of the haploid budding yeast nucleus in interphase. **(A)** The 16 chromosomes were modelled as bead-and-spring chains with 30 nm beads each comprising 3.2 kb of DNA. The chains are confined into the nucleus (1 μm radius sphere) and beads corresponding to centromeres were constrained in a sphere of radius 150 nm attached to the nuclear sphere to mimic the attachment to Spindle Pole Body (SPB) mediated by microtubules. The rDNA was restrained in a region occupying 10% of the nuclear volume at the opposite side of the nucleus with respect to the SPB. The telomeres were attracted to the nuclear envelope to have higher propensity to occupy the nuclear periphery, which is defined as the spherical shell, that is the closest to the nuclear envelope and occupies one third of the total volume of the nucleus (Methods). **(B)** The chromosomal polymer models, representing the genome-wide chromosome arrangement, were initialized as cylindrical solenoids of radius 150 nm. Next, the restraints on centromeres, rDNA, and telomeric particles were satisfied using a short preliminary run of Langevin dynamics, spanning 60τ_LJ_ (Methods). Finally, the system was relaxed with a 30,000τ_LJ_ run of Langevin dynamics, in which all the spatial restraints are in place. This run allows obtaining 10 steady state conformations per trajectory (one every 3,000τ_LJ_). Each strain was modelled in 1,000 independent replicates to obtain 10,000 genome-wide conformations per strain.

As an orthogonal validation of our model we compared the probability of contact among all chromosomal particles in the wild-type models with the experimentally measured intra- and inter-chromosomal contact frequencies observed by a 3C-derived technique [37,38]. In addition, we compared the predicted median telomere-telomere distances from our models with analogous experimental data obtained using imaging [39]. In both comparisons, we found that our models, based on the physical properties of chromatin and minimal biological restraints, accurately described the wild-type yeast nuclear organization (Supplementary Figures 1-2).

To determine if our computational models reproduce the experimentally measured low gene expression at the nuclear periphery, the predicted gene position relative to the nuclear periphery was correlated with genome-wide mRNA levels obtained by RNA-seq. Genomic regions within 30 kb of the ends of wild type chromosomes are poorly expressed, consistent with previous reports [21] (Figure 2A,B). Importantly, lower expression was also correlated with gene peripheral localization, as predicted by polymer modelling (Figure 2B). Because most subtelomeric sequences are also restricted to the perinuclear region, the above analysis confounds the contributions of sequence proximity to chromosome ends (1D effect) and proximity to the nuclear periphery (3D effect) to steady-state mRNA levels. However, we found that, while distance to the telomere and predicted location in the nuclear periphery are correlated, they are imperfectly so (Figure 2C). Especially for genes with low expression, the amount of time the gene is predicted to spend in the nuclear periphery is more highly correlated with expression than distance to the telomere in both linear (correlation = −0.093) and log space (Figure 2D). Furthermore, in a linear model that predicts expression from both of the two variables, % peripheral is a slightly more important feature (Supplementary Table 1). These data suggest that localization to the periphery, and not only distance from the telomere, is partially responsible for low expression.

**Figure 2.**
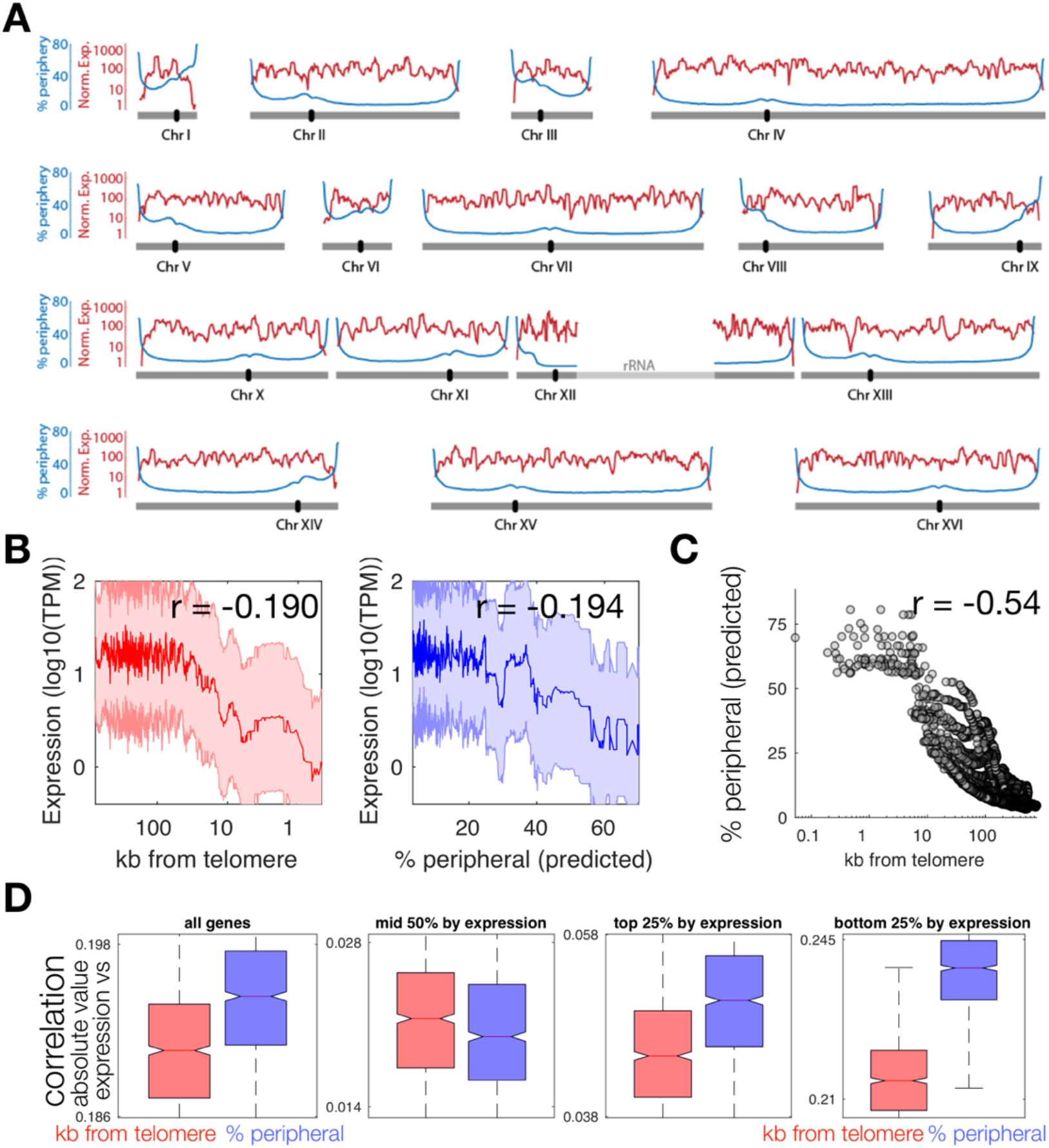
Localization in the nuclear periphery is associated with lower expression. **(A)** mRNA expression (red) and predicted time spent in the nuclear periphery (blue) are shown for each chromatin bead along each of the 16 yeast chromosomes. **(B)** Median expression level for genes binned by distance to the telomere (red) or by predicted % peripheral (blue). Correlation values are for Pearson correlation on unbinned data. **(C)** Predicted % peripheral is not perfectly correlated with distance from the telomere. **(D)** Gene expression is more strongly correlated with predicted % peripheral (blue) than with log(distance to the telomere). The effect is particularly strong for the most genes in the bottom quartile of expression. Boxplots show median correlation across 1000 random samplings of 90% of genes.

### Computational modelling and cell imaging validate nuclear reorganization after chromosomal rearrangements

To experimentally determine if spatial organization affects expression we next examined how large-scale chromosome rearrangements affect nuclear reorganization. In previously described Fusion Chromosome (FC) strains, up to three "donor" chromosomes were sequentially fused to the end of an intact "recipient" chromosome [29,30]. Centromeres were simultaneously removed from donor chromosomes to avoid formation of toxic dicentrics; telomere elements at the site of the fusion were also removed. Thus, like normal chromosomes, FCs contain two telomeres and one centromere (Figure 3A-B). These chromosome fusions only minimally changed the genomic content relative to wild type strains, since only 5 to 26 subtelomeric ORFs are lost during the fusion procedure (Supplementary Table S2). However, we hypothesized that FC strains would display dramatically altered interphase chromosome organization. Indeed, this is dependent on chromosome number and length, centromere attachment to spindle pole bodies, and telomere anchoring to the nuclear envelope (NE), all of which are altered in FC strains. Importantly, chromosome fusions led to a reduction in chromosome and centrosome number from 16 to 13, reduction of telomere number from 32 to 26, and lengthening of the longest chromosome arm (excluding chromosome XII, containing the variable rDNA array) from 1 to almost 4 Mbp (Figure 3B).

**Figure 3.**
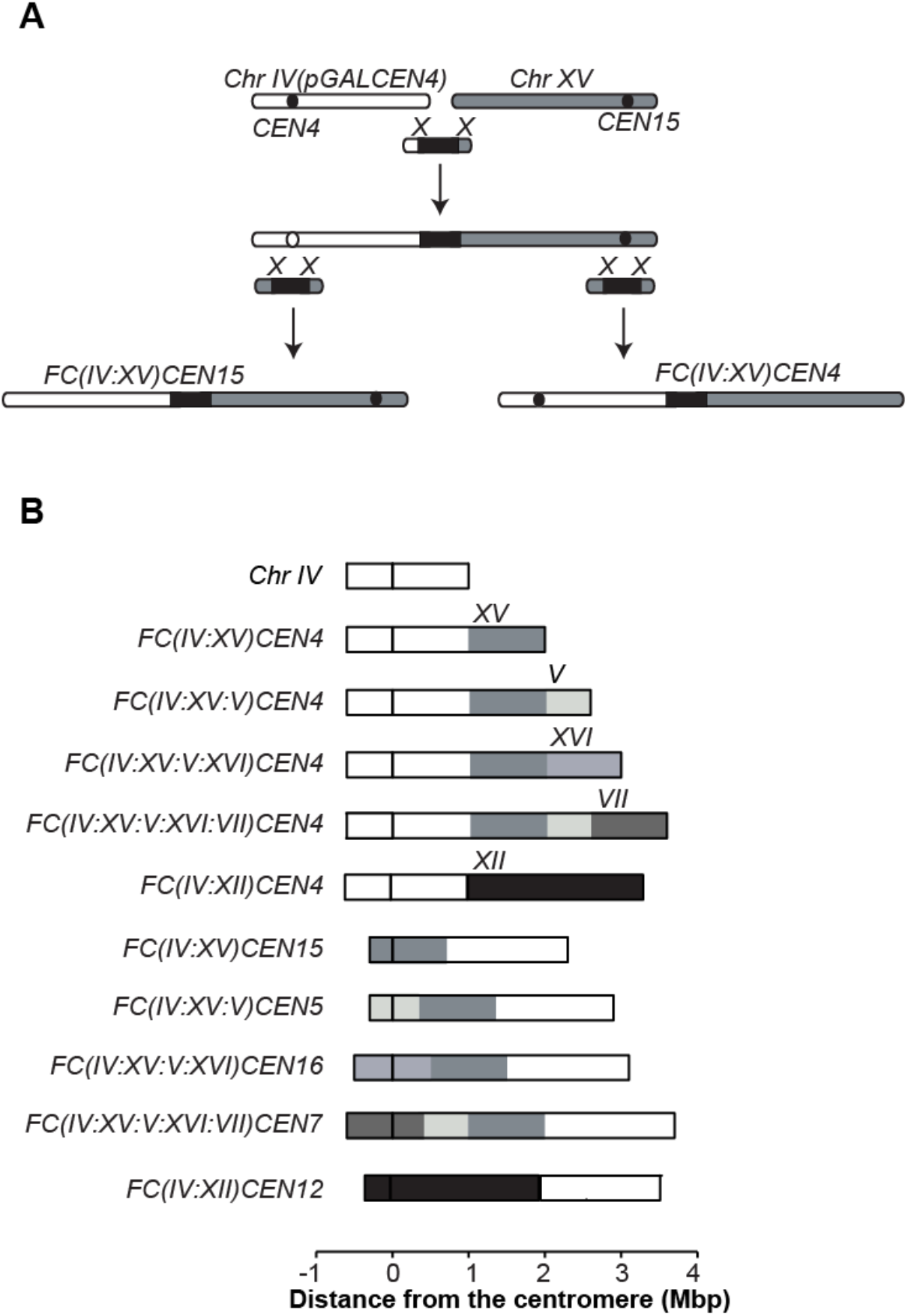
Generation of fused chromosomes strains. **(A)** The generation of fusion chromosomes (originally described in Neurohr et al. 2011 and Titos et al. 2014) starts with the integration of *pGAL1* sequence upstream of the centromere to be inactivated. Next, the chromosomes are fused by homologous recombination between a bridging PCR fragment and the telomeres of the chromosomes. Finally, the deletion of one of two centromeres and the excision of the *pGAL1* sequence, as appropriate, generates the *FC* strain. Black circle is the centromere, black rectangle is the selection marker. **(B)** Schemes of all the *FC* strains used in this work. Chromosome IV is shown for comparison.

We then applied the principles used in modelling wild-type nuclei to determine nuclear organization in the ten different FC strains (Figure 3A-B). Fusion chromosomes used in this study are named using the following convention: “FC” is followed by the chromosomes that comprise the fusion indicated in brackets, followed by the centromere of the recipient chromosome. Thus, strain *FC(IV:XV:V)CEN4* bears a fusion chromosome in which chromosome IV is the recipient, and chromosomes XV and V are the donors.

The model predicts two major changes in the FC strains. Firstly, large (> 300 nm) displacements of “donor” chromosomes away from the spindle pole body and slight (10-20 nm) displacement of “recipient” chromosomes towards the SPB (Figure 4 for IV:XII fusions, and Supplementary Figure 3 for all FCs). Secondly, the model predicts displacement of loci in the fused chromosomes away from the nuclear periphery. To quantify this non-intuitive prediction, we computed the distance from the nuclear periphery of all 10-kb loci from the surface of the nuclear sphere for all chromosomes in all strains relative to wild type. The model predicts that only loci on fused chromosomes are displaced away from the nuclear periphery, while the relative location of loci in non-fused chromosomes never varies by more than 50 nm (Figure 5A). Loci with the largest predicted displacement are located near the ends of fused chromosomes (Figure 5B).

**Figure 4.**
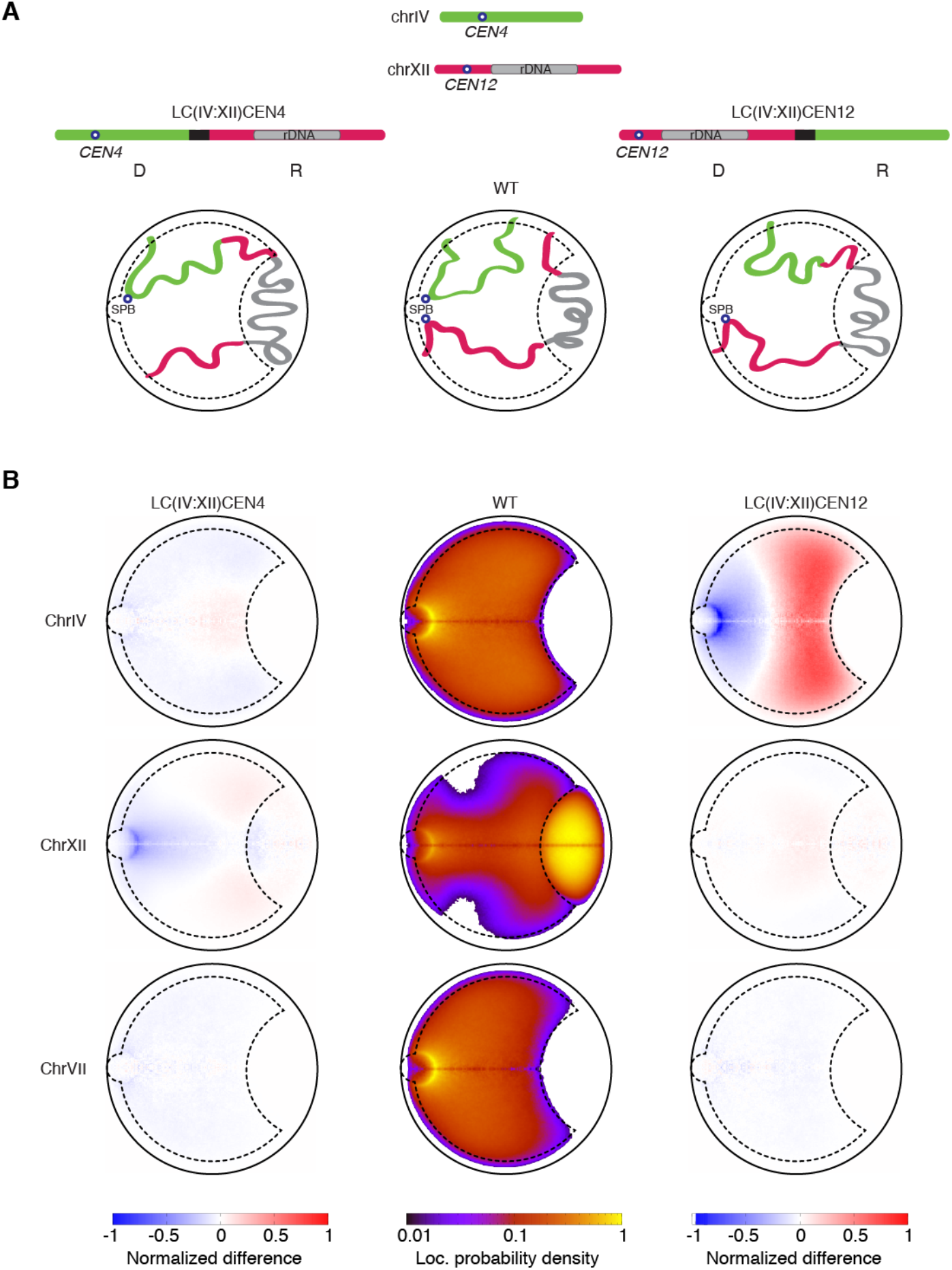
The donor chromosomes are predicted to be strongly displaced in the nucleus. **A.** Cartoon representations of the wild type, *FC(IV:XII)CEN4* and *FC(IV:XII)CEN12* strains. “Donor” and “recipient” chromosomes are labelled “D” and “R”, respectively. **B.** Predicted chromosome location probability densities for chromosomes IV, XII and VII in the wild-type strain (central column) and the *FC* strains *FC(IV:XII)CEN4* (left column) and *FC(IV:XII)CEN12* (right column), shown normalized by the wild type. The heat-maps show large differences in the positioning of the recipient and donor chromosomes, and almost no-difference in the nuclear organization of the largest non-fused one, chromosome VII.

**Figure 5.**
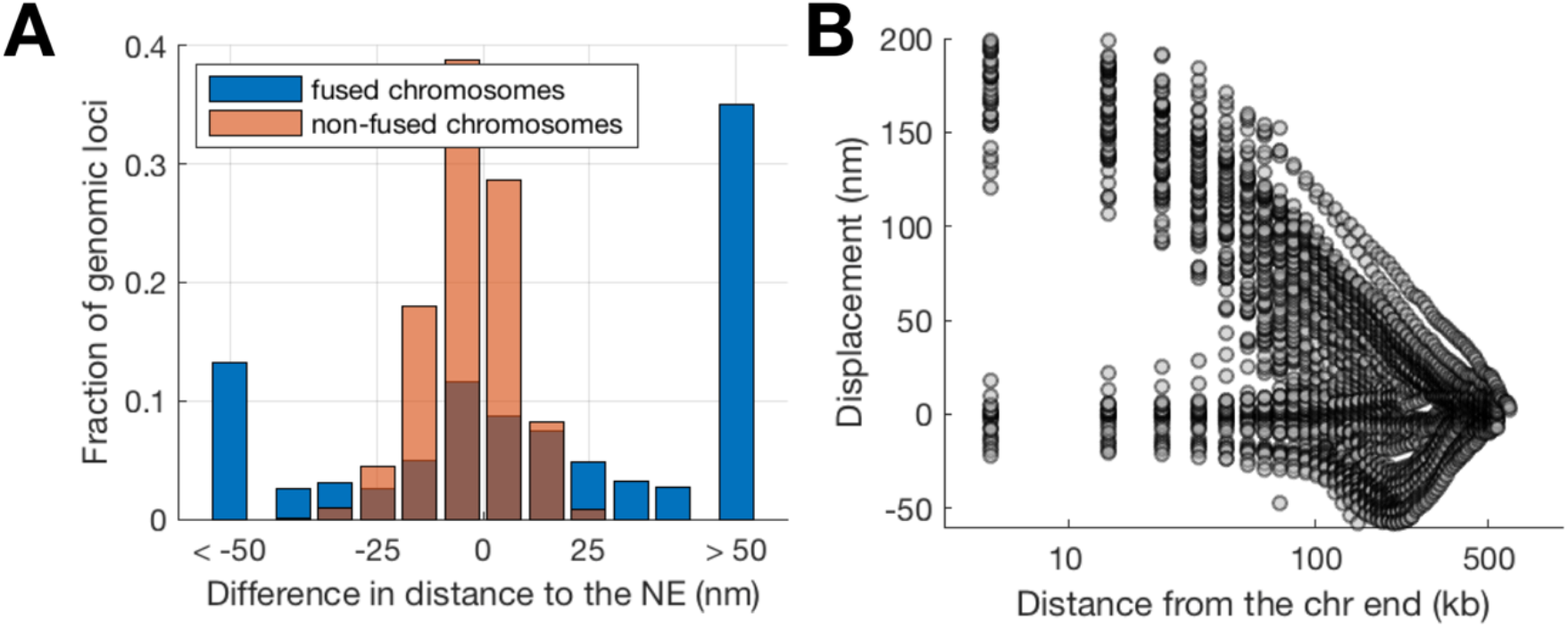
Loci near the end of fused chromosomes are predicted to be displaced away from the nuclear periphery. **(A)** The predicted displacement with respect to the nuclear envelope for loci in fused (blue) and non-fused (orange) chromosomes. **(B)** The predicted displacement with respect to the nuclear envelope for loci as a function of distance from the end of the chromosome in wild-type cells. Loci that are closer to the ends of the chromosomes exhibit a greater change away from the nuclear envelope.

To validate predicted chromosome displacement in FC strains we determined the distance of chromosome loci to each other, to the spindle pole body, and to the nuclear periphery using fluorescence microscopy in wild-type and FC strains. Loci in chromosome IV were visualized through TetR-mRFP and LacI-GFP reporters in cells bearing tetracycline and lactose operator arrays. These arrays were inserted respectively at the *TRP1* locus 10 kb from *CEN4* in the right arm of chromosome IV and at the *LYS4* locus in the middle of chromosome IV right arm, 470 kb away from *TRP1* (see scheme in Figure 6A). Distances were determined by live cell fluorescence microscopy in G1 cells expressing Spc42-GFP and Nup49-mCherry to label spindle pole bodies and the nuclear periphery, respectively. We found that the *CEN4*-associated *TRP1* locus is located in the vicinity of the SPB in wild-type and *FC(IV:XII)CEN4* nuclei (Figure 6B-D), whereas the same locus is displaced away from the SPB in *FC(IV:XII)CEN12* (Figure 6E). This agrees with model predictions that “donor” chromosomes are displaced away from the SPB, as compared to the wild-type configuration. Neither *TRP1* nor *LYS4* changed their distances from the nuclear periphery in either FC, consistent with model predictions. However, immunofluorescence and fluorescent in situ hybridization (IF-FISH) established that the *TEL4R*-proximal locus was closely associated with the nuclear periphery (labelled with a NPC-specific antibody) in wild type cells, whereas the mean distance between *TEL4R* and the nuclear periphery was increased in both *FC(IV:XII)CEN4* and *FC(IV:XII)CEN12* fusions (Figure 6F). Because all FC strains are derivatives of one of these two fusions, the *TEL4R* region is most likely displaced in these strains as well. This confirmed the model’s prediction that subtelomeric loci engaged in a chromosome fusion event are displaced away from the periphery (Figure 5A-B). Together, these results quantitatively confirm the model predictions that chromosome fusions lead to large changes in the sub-nuclear distribution of chromosome regions relative to both the spindle pole bodies and the nuclear periphery.

**Figure 6.**
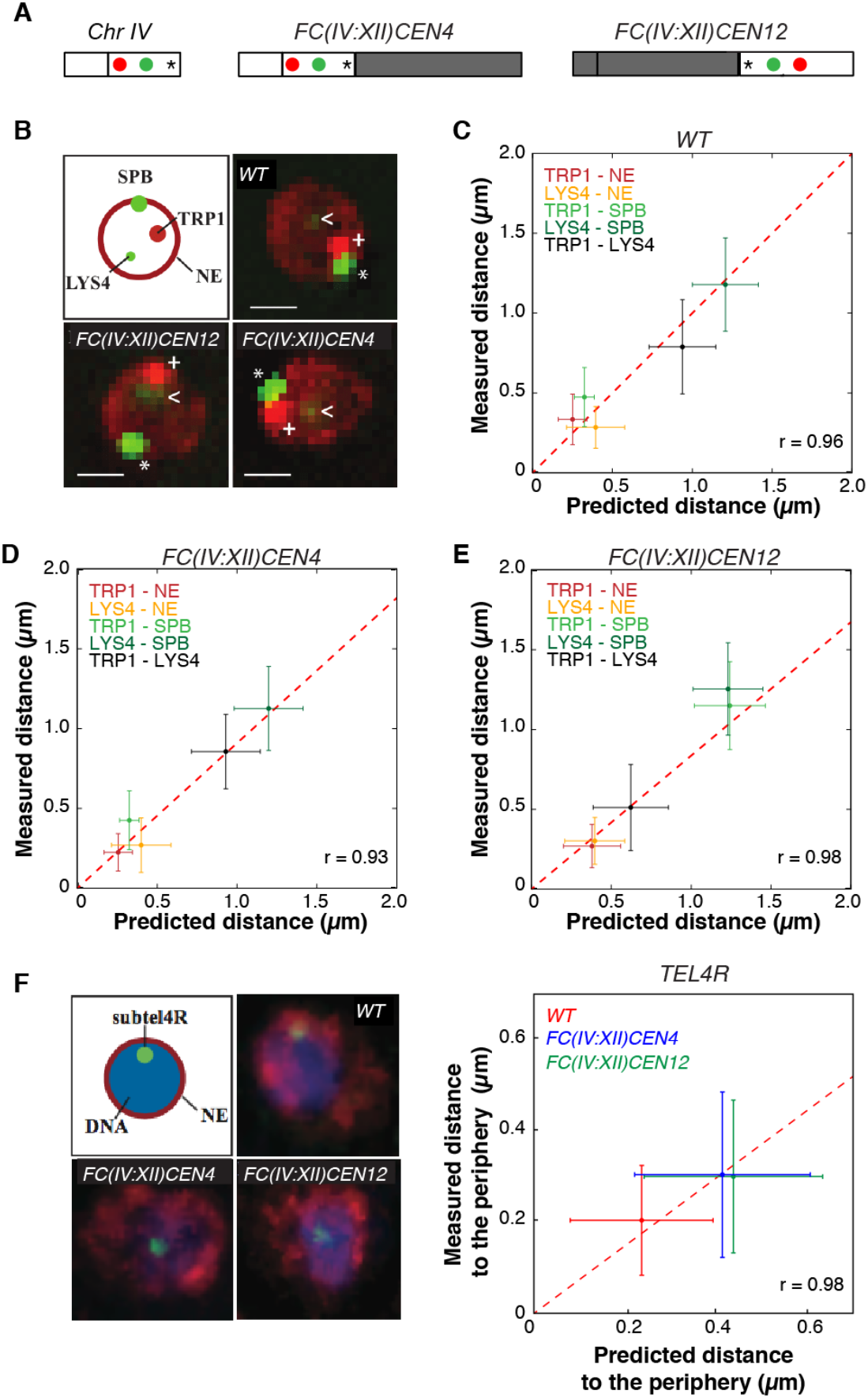
Validation of the polymer models by live and fixed cell microscopy. **(A)** Position of *TRP1* (red), *LYS4* (green), and *TEL4R* (asterisk) on chromosome 4 and its indicated FC derivatives. **(B)** Live cell microscopy of G1 cells of the indicated strains showing the localization of *TRP1* (red dot, marked with +), *LYS4* (faint green dot, arrowhead), the SPB (bright green dot, marked with an asterisk) and the nuclear periphery (red). **(C-E)** Correlation of measured and predicted distances between the indicated nuclear loci, the SPB and the nuclear periphery, in the indicated strains. **(F)** Fluorescence in situ hybridization and immno-fluorescence of G1 cells of the indicated strains showing the localization of *TEL4R* (green dot) and the nuclear periphery (NE) (red). Graphs show the mean and standard deviations from 50 cells / strain in C-E, 52 cells / strain in F, and 10.000 independent simulations in C-F. Scale bar, 1μm.

### Chromosomal rearrangements correlate with increase in expression of subtelomeric genes displaced away from the nuclear periphery

We next asked whether the genome reorganization caused by chromosome fusions led to changes in gene expression. We performed RNA-seq in the ten FC strains, with four independent RNA-seq replicate experiments per strain (Figure 3). Consistent with all FC strains having wild-type growth rates [29,30], global gene expression is not perturbed (Supplementary Figure 4). Thus, changes in gene location relative to the SPB and to other chromosomes did not affect expression. Further, we observe no relation between change in expression and distance to the lost or displaced centromere, suggesting that heterochromatin spreading around centromeres does not measurably affect expression (Supplementary Figure 5).

We then asked whether changes in expression correlated with changes in predicted gene position relative to the nuclear periphery. To obtain a more accurate value for expression in the absence of changes in nuclear location, for each gene we use the average expression level of that gene across all strains in which the percent peripheral is not predicted to increase or decrease by more than 1%. From this baseline expression value, we compared the fold change in expression for each strain with the predicted change in the amount of time that each gene spends in the nuclear periphery. Genes deleted during the fusion events were not considered. The results of this analysis show mild but statistically significant genome-wide expression changes for genes that change location relative to the nuclear periphery after chromosomal fusions (Figure 7A). A 10% increase in the amount of predicted time a gene spends outside of the nuclear periphery results in a ~10% increase in expression (Figure 7B-C). While effect on expression is weak, it is consistent across changes in localization and strains (Figure 7B-C) and remains if we limit our analysis to genes not involved in the stress response or only highly expressed genes (Supplementary Figure 6).

**Figure 7.**
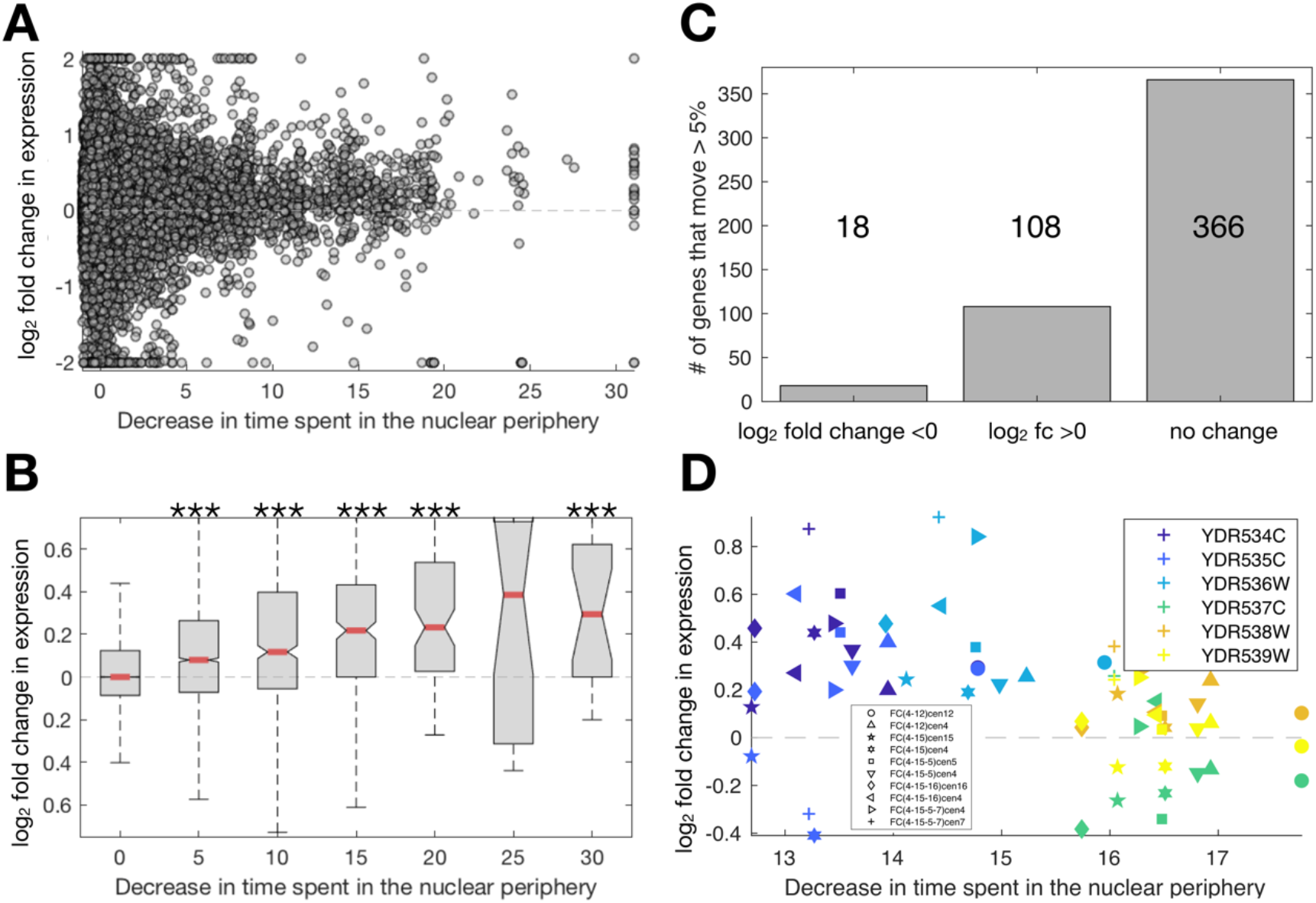
Gene displacement away from the nuclear periphery correlates with increased expression. **(A)** Shown for all genes and all strains are the fold change in expression and change in the predicted localization to the nuclear periphery. **(B)** The same data as in (A), with genes grouped by the predicted decrease in time spent in the periphery. Compared to genes whose localization does not change, groups of genes with significantly different changes in expression are marked with *** for when the p-value is <0.01 for a Kruskal-Wallis test with Tukey-Kramer multiple comparison correction. **(C)** The total number of genes that exhibit significant changes in expression due to changes in location. **(D)** Measured expression and predicted change in time spent in the nuclear periphery for the six genes around TEL4R. Colors mark genes, and symbols mark strains. This region is predicted to spend ~15% less time in the nuclear periphery, and all genes save YDR537C increase in expression.

Notably, genes with changes in both expression and localization were concentrated around subtelomeric regions of chromosomes engaged in fusion events, which models predict are the regions undergoing major displacement in *FC* strains. Examples of correlated changes in expression and localization are shown for the *TEL4R*-proximal region, which is perinuclear in wild-type cells but is displaced away from the nuclear periphery in *FC(IV:XII)*, and presumably in all other *FC* strains, as they are derivatives of *FC(IV:XII)* (Figure 3). Most genes in the *TEL4R*-proximal region show increased expression after displacement towards the nuclear interior (Figure 7D). For genes closer than 30 kb to the former telomere, these differences in expression correlate better with their distance to the deleted telomere than with their predicted time in the nuclear periphery, suggesting they are due to removal of telomere-associated silencing (Supplementary Figure 7). However, the predicted nuclear localization was a better predictor for expression level for genes located further than 30 kb from the former telomere, in both FC (Supplementary Figure 7) and wild-type cells (Supplementary Figure 8). Thus, distance from the nuclear periphery may dominate over distance to the telomere in determining the expression level of genes near chromosome ends that are located more than 30 kb away from telomeres.

## DISCUSSION

Interphase yeast chromosomes are organized with centromeres clustering around the spindle pole body (SPB), telomeres associating with the nuclear envelope (NE), and chromosome arms extending between these two anchoring points in a brush-like fashion. How this organization affects nuclear functions is not fully understood. Previous studies reported altered expression of subtelomeric genes in mutants that disrupt heterochromatin formation or telomere clustering [19,21]. Importantly, these studies did not directly address the role of three-dimensional chromosome organization, as the genetic perturbations used (depletion of histone H4, and mutations of the silencing factor *SIR3* and of the telomere tethering proteins *YKU70* and *ESC1*) affected multiple processes, including heterochromatin formation, genome-wide gene expression and DNA repair. In this study, we used tailored chromosome fusions (FC cells) to alter interphase nuclear organization in otherwise wild-type cells. Computational modelling validated with single cell imaging revealed significant changes in nuclear organization after these chromosome fusion events. This highlights the power of polymer-based modelling approaches to reproduce chromosome-level organization of wild type yeast nuclei, and to predict the reorganization caused by chromosome rearrangements, based only on minimal imposed constraints.

Our analysis reveals that genome-wide gene expression levels remained generally unaffected by changes in chromosome organization. However, we also find that chromosome fusions result in consistent and reproducible increases in expression, with over 100 subtelomeric genes exhibiting a mild but significant increase. This is consistent with normal growth of FC strains in rich media [30], and with recent reports that overall transcription is not affected by fusion of all yeast chromosomes into one or two mega-chromosomes [40,41]. These two studies also reported de-repression of subtelomeric genes near chromosome fusion sites, which was attributed to disruption of telomeric silencing. These studies used one to three RNA-seq biological replicates, whereas by combining analysis of mRNA-seq data across multiple FC strains, we effectively have 38 biological replicates. Accurate quantification of expression changes of less than 50% requires more than 10 replicates [42], potentially explaining why we identify a relatively higher number of genes with changes in expression of 10 to 20%. Because increased expression of these genes is correlated with both their 1D distance to the former telomere and their 3D distance to the periphery, both telomere-associated silencing and spatial displacement away from the nuclear periphery may contribute to increased expression levels of subtelomeric genes. Our results suggest that removal of telomere-associated silencing is likely to be the dominant factor affecting genes <30 kb from the former telomere, whereas 3D distance to the periphery may be the major factor affecting genes further away. Interestingly, distinct histone deacetylases target these two subtelomeric regions: Sir2 and Hda1 prevail within 30 kb of telomeres, whereas Rpd3 targets regions 30 to 50 kb away from telomeres [43,44]. Displacement from the nuclear periphery may alter chromatin organization in these regions leading to changes in expression.

It is interesting to consider our results in the context of previous studies on the mechanisms of subtelomeric silencing. Transcription levels are known to decrease in proximity to telomeres (reviewed in [45]). Moreover, gene targeting to the nuclear periphery, either by integration of reporters in subtelomeric regions or by artificial anchoring to perinuclear proteins, leads to silencing that is dependent on perinuclear enrichment of SIR factors [16-19]. These observations led to the hypothesis that the nuclear envelope is a transcriptionally repressive environment due to the local accumulation of repressive factors. However, a truncated telomere that does not localize to the nuclear periphery can still support silencing of a *URA3* reporter [46], and microarray analysis showed that almost 80% of subtelomeric genes were still silenced after telomere detachment form the nuclear periphery in *esc1 yku70* mutants [19]. These findings raised the possibility that subtelomeric gene position and expression are independent from each other. In contrast, our results suggest that displacement from the nuclear periphery affects the expression levels of native subtelomeric genes, but that this effect is relatively mild, which may have escaped previous analysis using growth on selective media or microarrays. These findings support the hypothesis that regulation of perinuclear localization of subtelomeric genes (e.g. by telomere detachment) may affect their expression in response to environmental signals. Since chromosome detachment in FC strains examined here caused relatively mild changes in expression, it remains unclear to what extent changes in position may contribute to induction of subtelomeric gene expression in stress conditions.

In summary, the data presented here establish FC strains as an excellent tool to study the relationship between nuclear organization and function without resorting to mutations, which may have unintended effects, and suggest that chromosome position plays a role in setting gene expression levels for subtelomeric genes. Similar approaches could be used to test to what extent other nuclear processes that are sensitive to gene position, such as DNA replication timing and DNA damage repair [47,48] are also affected by global changes in nuclear organization.

## MATERIALS AND METHODS

### Polymer modelling

Each yeast chromosome of wild type and fused chromosome strains was modelled using a bead-and-spring polymer model previously used and validated for modelling chromatin fibers [34]. This model consists of 3 different energy contributions each describing a general physical property of the chain:

1. *Excluded volume (Purely repulsive Lennard-Jones potential).* Each particle occupies a spherical volume of diameter equal to 30nm and cannot overlap with any other particle in the system. Considering the typical compaction ratio of the chromatin fiber in yeast [49,50], each particle contains about 3.2 kb of DNA.
2. *Chain connectivity (Finite extensible nonlinear elastic potential).* Consecutive particles on the chain are connected with an elastic energy, which allows a maximum bond extension of 45 nm. The simultaneous action of the excluded volume and the chain connectivity prevents chain crossings.
3. *Bending rigidity (Kratky-Porod potential).* The bending properties of an ensemble of polymer chains is usually described in terms of the *persistence length*, which is the length-scale where the chain changes its behaviour from rigid to flexible. According to the bending properties experimentally measured for the yeast chromatin fiber [49,51,52], the persistence length of each model chain was set to 61.7 nm for internal regions of the chromosomes, and to 195.0 nm for the terminal ones. The regions of the chains corresponding to the telomeres (the 20 kb at the chromosomes ends), in fact, are more compact and rigid [53].

Since the modelling aims to describe the chromosomal configuration of haploid strains, the total number of beads in the system is 4,062, resulting from the presence of one copy of each yeast chromosome (Supplementary Tables S3-S4). Each chromosome is initially folded in a solenoidal arrangement, where a rosette pattern is repeatedly stacked to yield an overall linear, rod-like conformation, see Figure 1 [34,35,54].

The chromosome chains are consecutively placed inside a sphere of radius 1.0 centred in the origin (0,0,0). This sphere describing the typical shape of the yeast nucleus in G1, according to imaging data, interacts with the chromosome particles as a rigid wall. To obtain the initial chromosome nuclear locations, the position of the chromosome centres is picked in a random, uniform way inside the nucleus, and the orientation of the rod axis is chosen randomly. The iterative placement proceeds from the longest to the shortest chromosome in a way that the newly added chromosomes must not clash with previously placed ones. In case of a clash, the placement attempt is repeated. Next, the following biological restraints (i-iii) are satisfied using a short preliminary run of Langevin dynamics, spanning 60τ_LJ_, where τLJ is the Lennard-Jones time and is used as the time unit in LAMMPS:

i. To simulate the tethering of the centromeres to the spindle pole body (SPB), the motion of the centromeres particles was restrained into a spherical compartment of radius R_SPB_=150 nm centered in c_SPB_=(−850,0.0,0.0).
ii. rDNA particles were restrained to a region occupying 10% of the total nuclear volume and located at the opposite side of the SPB, to simulate the nucleolus. Nucleolar volume was derived from experimental measurements. This region was defined by the intersection of the nuclear sphere with a sphere of radius R_NUCL_=640.92nm whose center is located at c_NUCL_=(1000,0.0,0.0). Conversely, the other no-rDNA particles of the chromosome models were restrained to stay out of the same nucleolar compartment.
iii. Finally, to represent the tendency of the telomeres to stay anchored to the nuclear envelope (NE), the periphery of the sphere (a shell within R_PER_=126nm from the nuclear envelope which accounts for one third of the nuclear volume) was attracting for the terminal particles of the chromosome chains. This effect, unexplored so far, was accomplished using a Lennard-Jones attraction [55].

The restraints (i) to (iii) were imposed applying on each of the involved particles a force F, only when the particle did not satisfy the confinement conditions, using the option indent of the software LAMMPS [56]:

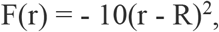

where r was the distance from the particle to the center of the sphere, and R was the radius of the sphere.

In the *FC* strains, the chromosomes involved in the fusion were attached to each other using additional connectivity bonds (points 2 above) between the telomeres involved in the fusion process. These telomeres, which were attracted to the periphery in the wild type strain models, behaved as internal chromosomal sequences in the *FCs* strains, and lost the telomeric attraction to the nuclear envelope.

Finally, the system was relaxed using a run of Langevin dynamics of 30,000_τLJ_, and one conformation every 3,000τ_LJ_ (10 models per trajectory) was retained for analysis. Replicating the complete simulation 1,000 times generated 10,000 genome-wide conformations per strain.

### Previously published modeling approaches

*S. cerevisiae* genome has been previously modelled using two main restraint-based approaches. First, Chromosome Conformation Capture datasets have been used as input restraints to reconstruct the 3D confirmation of the yeast genome [37,38,57]. Second, and in a similar approach used in our work, models were built using genome tethering to nuclear elements as restraints [32,33]. The differences between our approach and these previously published are minimal. For example, in this work, the genome was represented as series of spherical beads compared to cylinders previously used by [33]. Moreover, the initial conditions of the simulation, the confinement of the genome as well as the minimization protocols were different in our work compared to the used by [32]. However, these differences are likely to minimally change the final conclusions of our modelling approach compared to those previously published.

### Strains, cell growth and live cell microscopy

*Saccharomyces cerevisiae* strains are derivatives of S288c. TetO/LacO cells and chromosome fusions were previously described; briefly, haploid cells were transformed with a PCR fragment encoding a selection cassette flanked by sequences with homology to subtelomeric regions [29,30]. Live-cell microscopy was carried out with a Leica imaging system (AF6000). All live-cell images were acquired at 30°C with a ×100 objective lens. Eleven 0.2 μm thick z-sections were collected. Distances were measured between local maxima on single planes using ImageJ (http://rsb.info.nih.gov/ij/) and Microsoft Excel although for clarity, figures are represented as 2D maximum projections of whole-cell Z-stacks. Graphs and statistical analysis (*t*-test allowing for unequal variance) were performed with R and Excel (Microsoft).

### Immunofluorescence and fluorescence in situ hybridisation (IF-FISH)

To make FISH probes, a 6 kb PCR fragment in the *TEL4R* region was amplified from genomic DNA with primers: (5’-ATCTTTCCTTACACATAAACTGTCAAAGGAAGTAACCAGG-3’) and (5’-GTAACATACAAACTCAACGCCTACTAAGATTAATACATCA-3’), and labelled with Alexa Fluor 488 by nick translation using the FISH Tag-DNA Multicolor Kit (Invitrogen). FISH-IF was performed essentially as described [58], with minor modifications. Overnight cultures (1–2^10^ cells/ml) were treated with 10 mM DTT in 0.1 M EDTA/KOH, pH 8.0, treated with 0.4 mg/ml Zymolyase 100T (Seikagatu) for 15 min at 30°C in YPDA medium containing 1.1 M sorbitol (YPDA-S). This treatment allowed the cells not to be completely converted into spheroplasts, but partially retained their cell walls, to help stabilize their three-dimensional structure. Partially spheroblasted cells were fixed for 20 min with 3.7% paraformaldehyde in YPD-S at room temperature. Cells were recovered by centrifugation (1000 *g* for 5 min), washed three times in YPD-S, resuspended in YPDA, spotted on Teflon slides, left to air-dry for 5 min, then immersed in methanol for 6 min and in acetone for 30 s at 20°C. Slides were then rinsed in a phosphate-buffered saline containing 0.1% Triton X-100 (PBS-T) and 1% ovalbumin, incubated overnight at 4°C (or for 1 h at 37°C) with anti-Nuclear Pore Complex Proteins antibody [Mab414] - ChIP Grade (ab24609), diluted 1:2 in PBS-T. Slides were then washed in PBS-T and incubated with preabsorbed Cy5 AffiniPure Goat Anti-Mouse IgG (H+L) diluted to 0.025 mg/ml in PBS-T at 37°C for 1 h. Next, slides were fixed again in PBS containing 3.7% freshly paraformaldehyde for 20 min and incubated overnight in 4x SSC, 0.1% Tween 20, 20 μg/ml of RNase A at room temperature. Slides were then washed in water, sequentially immersed for 1 min in 70, 80, 90, and 100% ethanol at - 20°C, and air-dried. Slides were then denaturated at 72°C with of 70% formamide and 2 SSC, immersed for 1 min sequentially in 70, 80, 90, and 100% ethanol at −20°C and air-dried. The hybridization solution (50% formamide, 10% dextran sulfate, 2x SSC, 0.05 mg/ml labeled probe, and 0.2 mg/ml single-stranded salmon sperm DNA) was then applied and slides were incubated at 10 min at 72°C. Slides were incubated for 48 h at 37°C to allow probe hybridization, washed twice for 10 min each at 42°C in 0.05x SSC and twice in BT buffer (0.15 M NaHCO, 0.1% Tween 20, pH 7.5) with 0.05% BSA for 30 min. After three washes in BT buffer, slides were mounted in 1x PBS, 80% glycerol, 24 μg/ml 1,4diazabicyclo-2,2,2,octane, pH 7.5. Images from IF-FISH were acquired on a confocal microscope (Leica TCS SPE) with a ×100 objective.

### RNA-seq

Cells were harvested by centrifugation and RNA was extracted from fresh pellets using the RiboPure Yeast Kit (Ambion). RNA concentrations were determined using a NanoDrop 1000 (Thermo Scientific), while quality and integrity was checked using a Bioanalyzer 2100 (Agilent Technologies). RNA-seq was performed on a HiSeq2000 (Illumina). Paired-end reads of 50 bp were aligned to the reference *S. cerevisiae* genome (R64-1-1) using using kallisto quant -i orf_coding_all.idx -o output - b 100 read1_file.fastq.gz read2_file.fastq.gz.

To obtain a robust and accurate wild-type expression level for each gene, we averaged across strains. For each strain in which the gene is predicted to increase or decrease time spent in the nuclear periphery by less than 1% we took the median expression value across all strains (four independent RNA-seq replicate experiments per strain). Fold-change in expression was calculated as the log2 ratio of expression in the *FC* strain divided by expression in this median expression value. Similar results are obtained if expression for the wild-type control strain are used, but as many of the genes are expressed at very low levels, and hence represented by very few reads, averaging across strains is more robust to random counting noise.

## Data accessibility

Data and code are available at https://github.com/Lcarey/DiGiovanni_DiStefano_FC and RNA-seq raw data are available as GEO accession Nr GSE108261 at https://www.ncbi.nlm.nih.gov/geo/query/acc.cgi?acc=GSE108261

## Acknowledgements

We thank Guillaume Filion and all members of the Mendoza lab for comments, and the CRG Genomics Unit for assistance with RNA sequencing. M.M. acknowledges support of the European Research Council Starting Grant 2010-St-20091118 and the Spanish Ministry of Economy and Competitiveness BFU2012-37162. L.B.C. was supported by Agència de Gestió d’Ajuts Universitaris i de Recerca (2014 SGR 0974) and the Spanish Ministerio de Economía y Competitividad and FEDER (BFU2015-68351-P). M.A.M-R was supported by the European Research Council under the 7th Framework Program FP7/2007-2013 (ERC grant agreement 609989), the European Union’s Horizon 2020 research and innovation programme (grant agreement 676556) and the Spanish Ministry of Economy and Competitiveness (BFU2013-47736-P). We also knowledge support from ‘Centro de Excelencia Severo Ochoa 2013-2017’, SEV-2012-0208 to the CRG.

## Author Contributions

MAMR & MM conceived the project. FDG performed experiments. MDS and DB performed simulations. MDS and LBC analyzed the data. All authors contributed to the writing of the manuscript.

## SUPPLEMENTARY TABLES

**Supplementary Table 1.** Predicted % peripheral is a significant feature in a linear model that predicts mRNA expression in wild-type cells from using both zscored predicted % peripheral and zscored log distance to the telomere. Shown are linear models that predict expression from both features independently (bottom) or with an interaction term (top). In the non-interaction model, predicted % peripheral is a stronger predictor of expression than log(distance to the telomere). In the interaction model, the interaction term is the strongest predictor.

|  | <u>Estimate</u> | <u>SE</u> | <u>tStat</u> | <u>pValue</u> |
| --- | --- | --- | --- | --- |
| Dist2Tel_log | 0.056238 | 0.0156 | 3.6083 | 0.00031 |
| PercentPeripheral | -0.016564 | 0.0180 | -0.9225 | 0.35627 |
| Dist2Tel_log:PercentPeripheral | 0.04074 | 0.0052 | 7.7097 | 1.4558e-14 |

|  | <u>Estimate</u> | <u>SE</u> | <u>tStat</u> | <u>pValue</u> |
| --- | --- | --- | --- | --- |
| Dist2Tel_log | 0.0734 | 0.0155 | 4.7359 | 2.2288e-06 |
| PercentPeripheral | -0.0874 | 0.0155 | -5.6389 | 1.7857e-08 |

**Supplementary Table S2:**
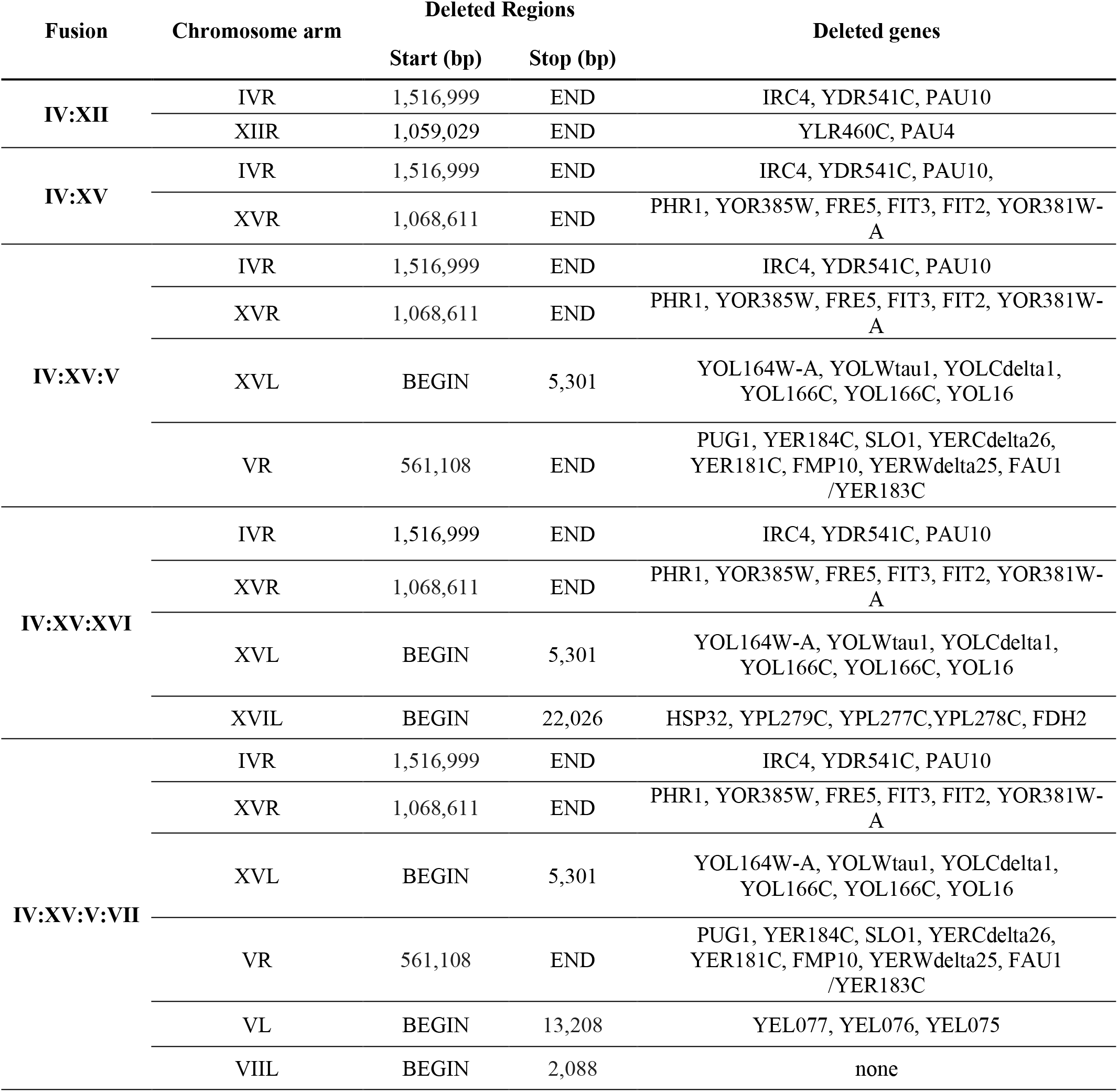
Deleted genes in *FC* strains

**Supplementary Table S3.** Parameters of the polymer models.

| Parameter | Value | Reference |
| --- | --- | --- |
| Number of chromosomes | 16 | 1 |
| Chromosome persistence length | 61.7 nm | 2 |
| Chromosome persistence length (last 30kb) | 195.0 nm | 3 |
| Nuclear diameter | 2 $\mu\text{m}$ | This study |
| Particle DNA content | 3.2 kb | 3 |
| Diameter of euchromatin segments | 30 nm | 3 |
| Number of repeats of the 9.1kb rDNA region | 102 | 1 |
| Chains can cross each other | No | 4 |

1. Cherry JM, et al. (1997) Genetic and physical maps of *Saccharomyces cerevisiae*. *Nature* 387 (6632 Suppl):67-73
2. Tjong et al 2012
3. Bystricky et al 2014
4. Rosa and Everaers 2008

**Supplementary Table S4.**
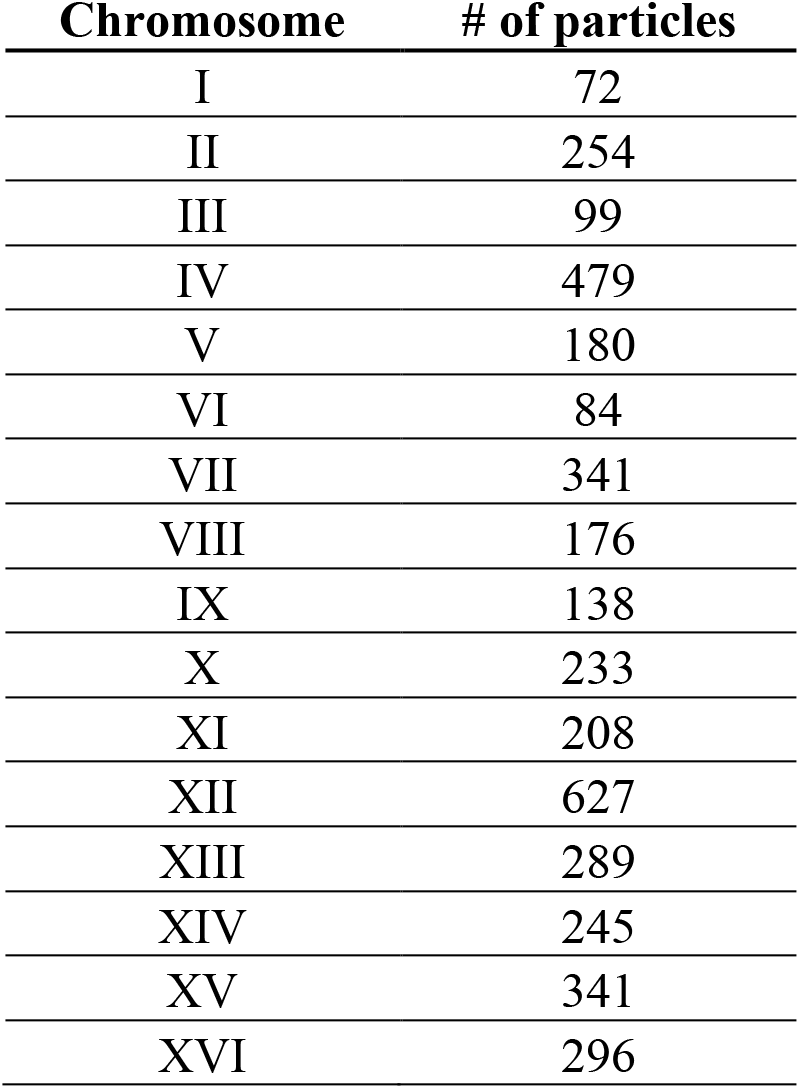
Number of particles per chromosome chain in the polymer models.

| **Chromosome** | **# of particles** |
| --- | --- |
| I | 72 |
| II | 254 |
| III | 99 |
| IV | 479 |
| V | 180 |
| VI | 84 |
| VII | 341 |
| VIII | 176 |
| IX | 138 |
| X | 233 |
| XI | 208 |
| XII | 627 |
| XIII | 289 |
| XIV | 245 |
| XV | 341 |
| XVI | 296 |

## SUPPLEMENTARY FIGURES

**Supplementary Figure 1.**
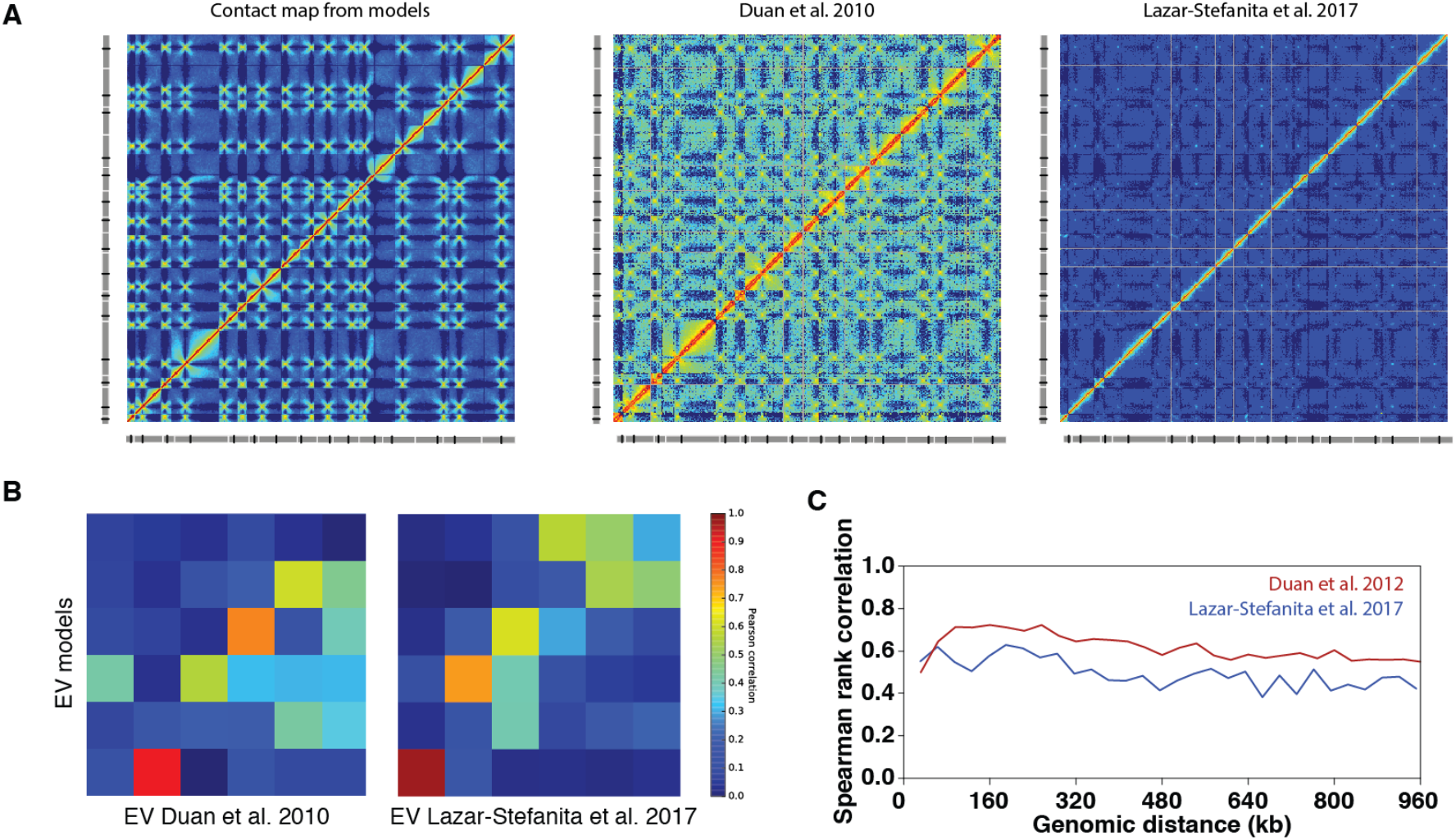
Comparison of the experimental and predicted contact maps. **(A)** DNA contact maps obtained from computational models and from experimental approaches. **(B)** Spectral decompositions of the maps show significant internal correlations up the first 6th eigenvalues [59]. **(C)** Correlations between elements grouped by genomic distance are significant up to the typical length of a budding yeast chromosome arm.

**Supplementary Figure 2.**
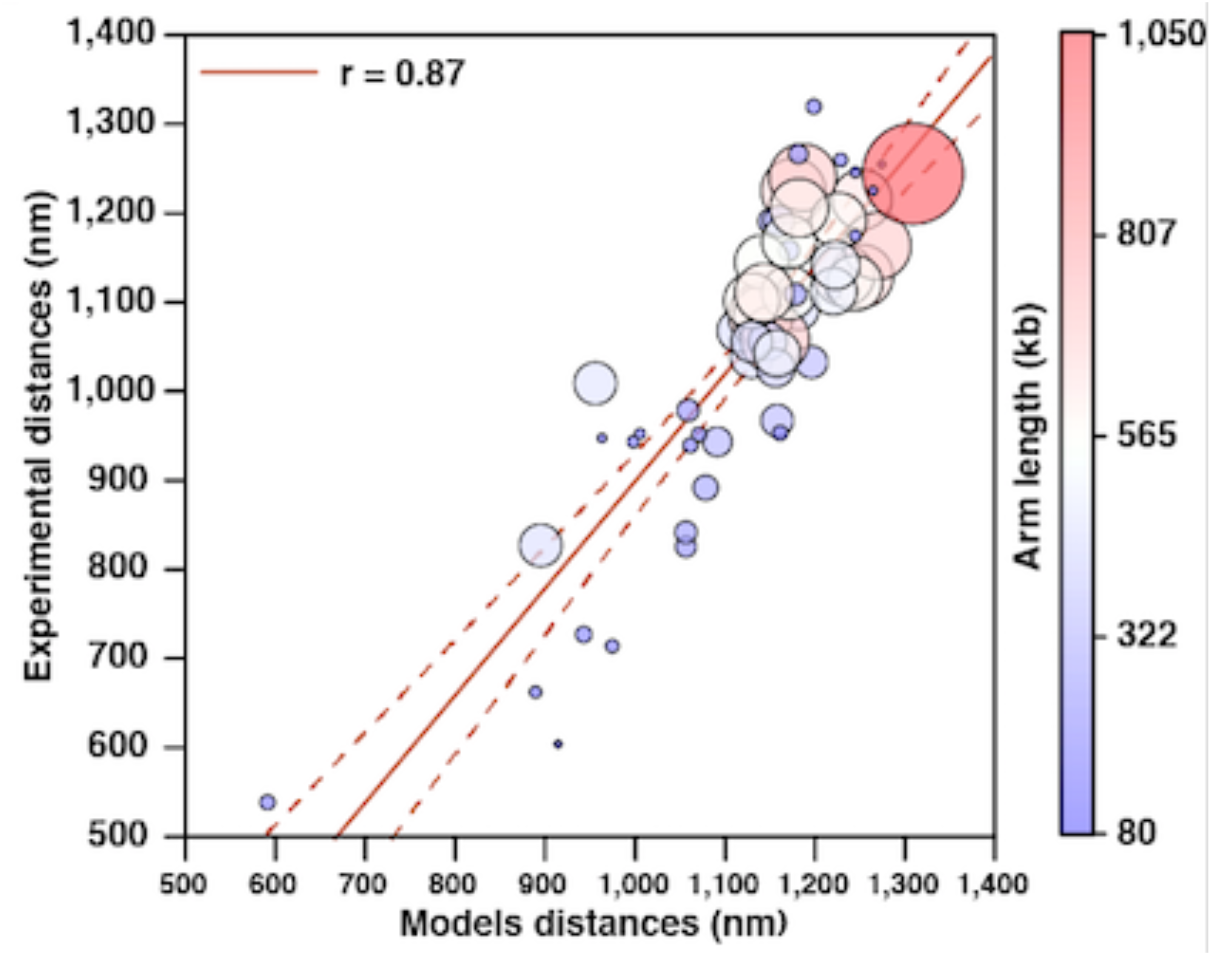
Correlation between measured median telomere-telomere distances in [39]. Analogous computations done on the wild type models show a significant agreement between models and experimental results.

**Supplementary Figure 3.**
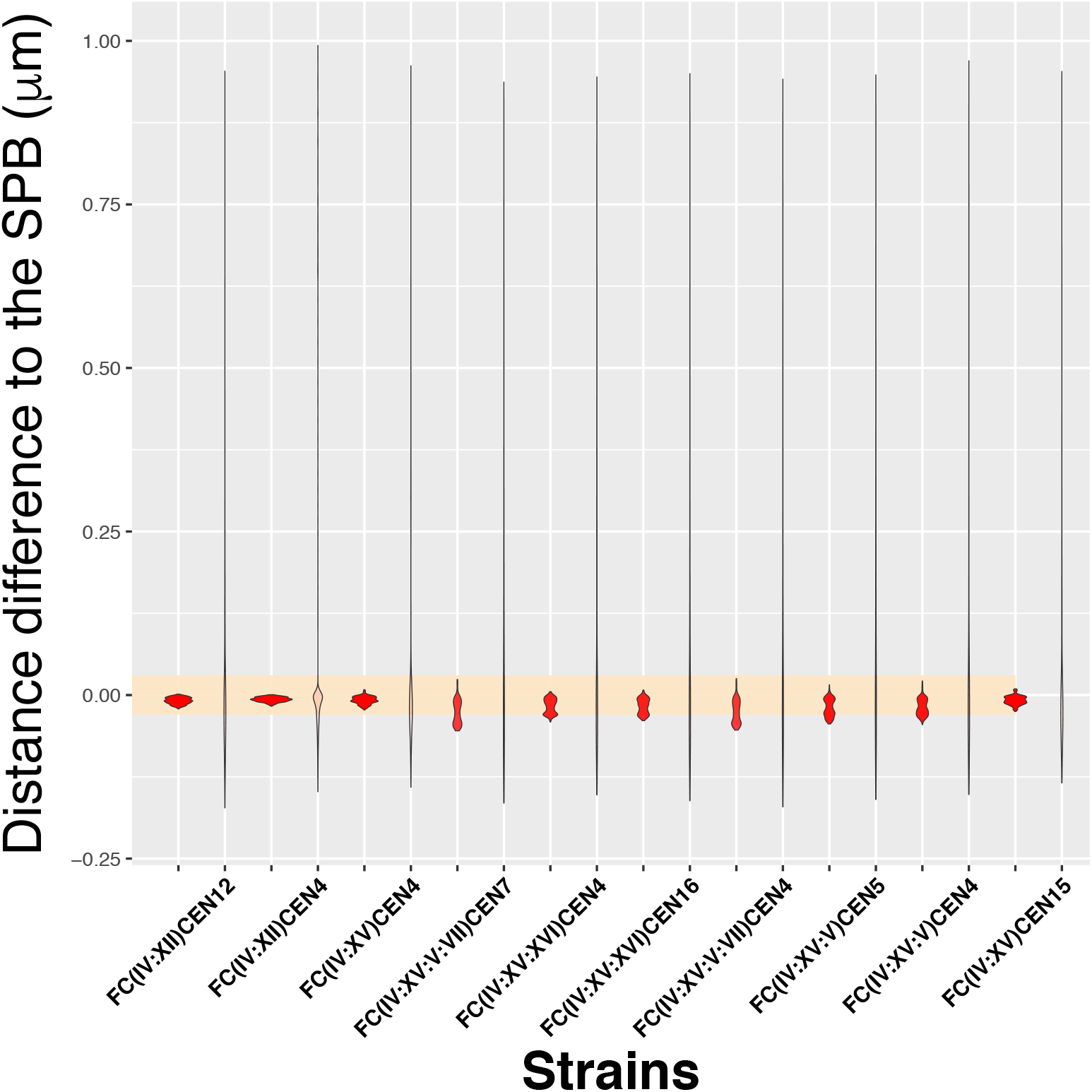
Displacement of fused chromosomes away from the SPB. Two distributions of the difference in average distance away from the SPB are shown for loci in *FC* strains with respect to the wildtype, including the non-fused chromosomes (left) and the fused chromosomes (right). The yellow reference area indicates the interval between −50 nm and 50 nm. Only fused “donor” chromosomes show displacements away from the SPB larger than 50 nm.

**Supplementary Figure 4.**
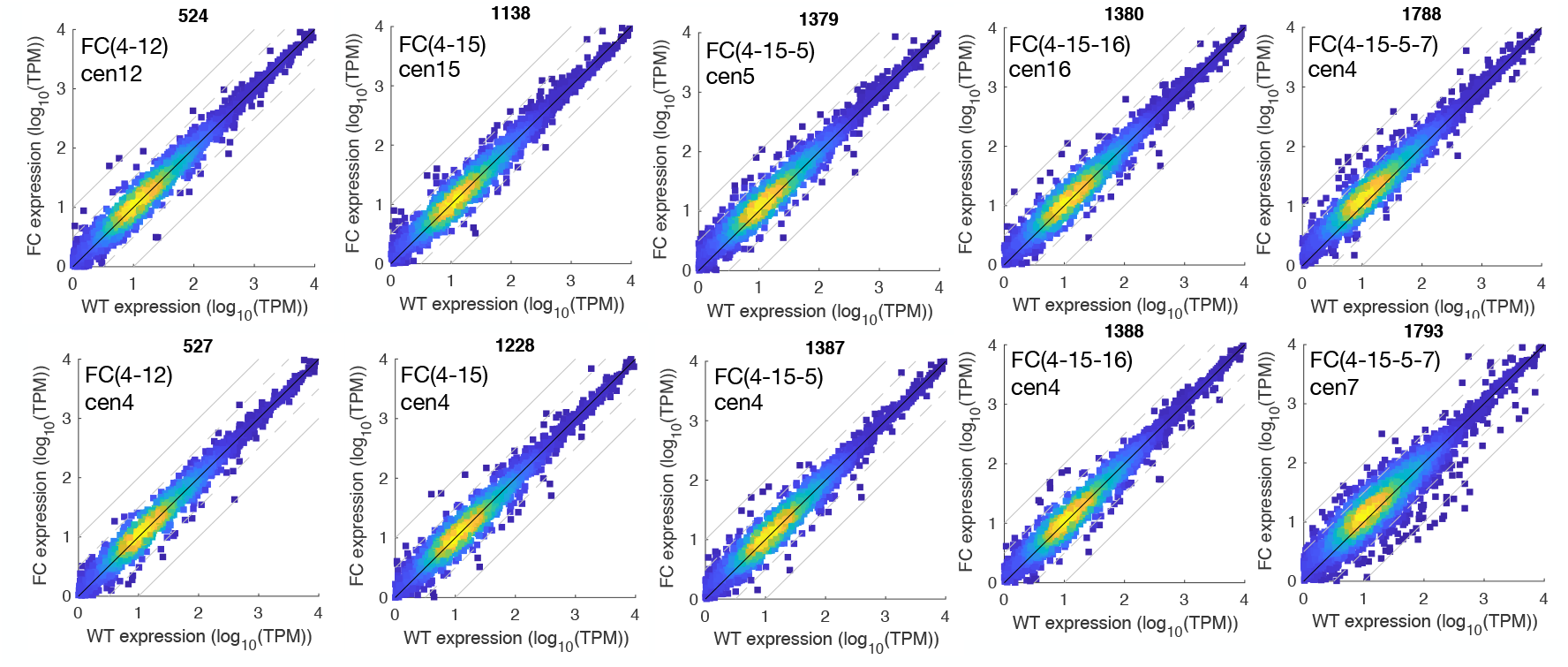
*FC* strains do not exhibit large-scale changes in gene expression. Expression levels in each *FC* strain are compared to expression in the WT-strain. Solid grey lines show a fold change of 1.0, dashed grey lines show a fold change of 0.5.

**Supplementary Figure 5.**
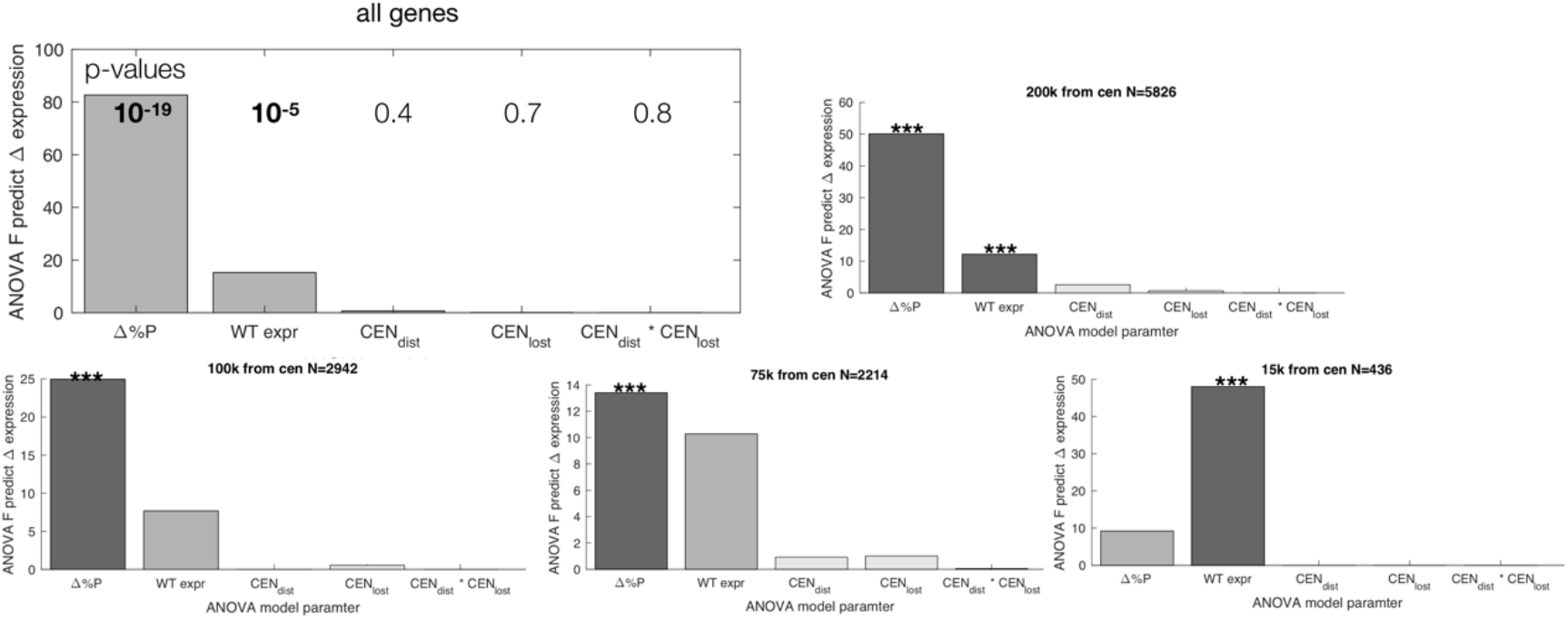
No significant effect of distance to the lost or displace centromere on changes in expression in the FC strains. ANOVA F-Statistics for measured change in expression (log2(FC/WT)) for all genes, or only genes 200kb, 100kb, 75kb or 15kb from the centromere for multivariate ANOVA in a model shows that the only significant predictors of change in expression are the predicted change in the time the genes spends in the nuclear periphery and, to a lesser extent, the expression of that gene in WT cells (*** = p<0.05 after multiple hypothesis testing).

**Supplementary Figure 6.**
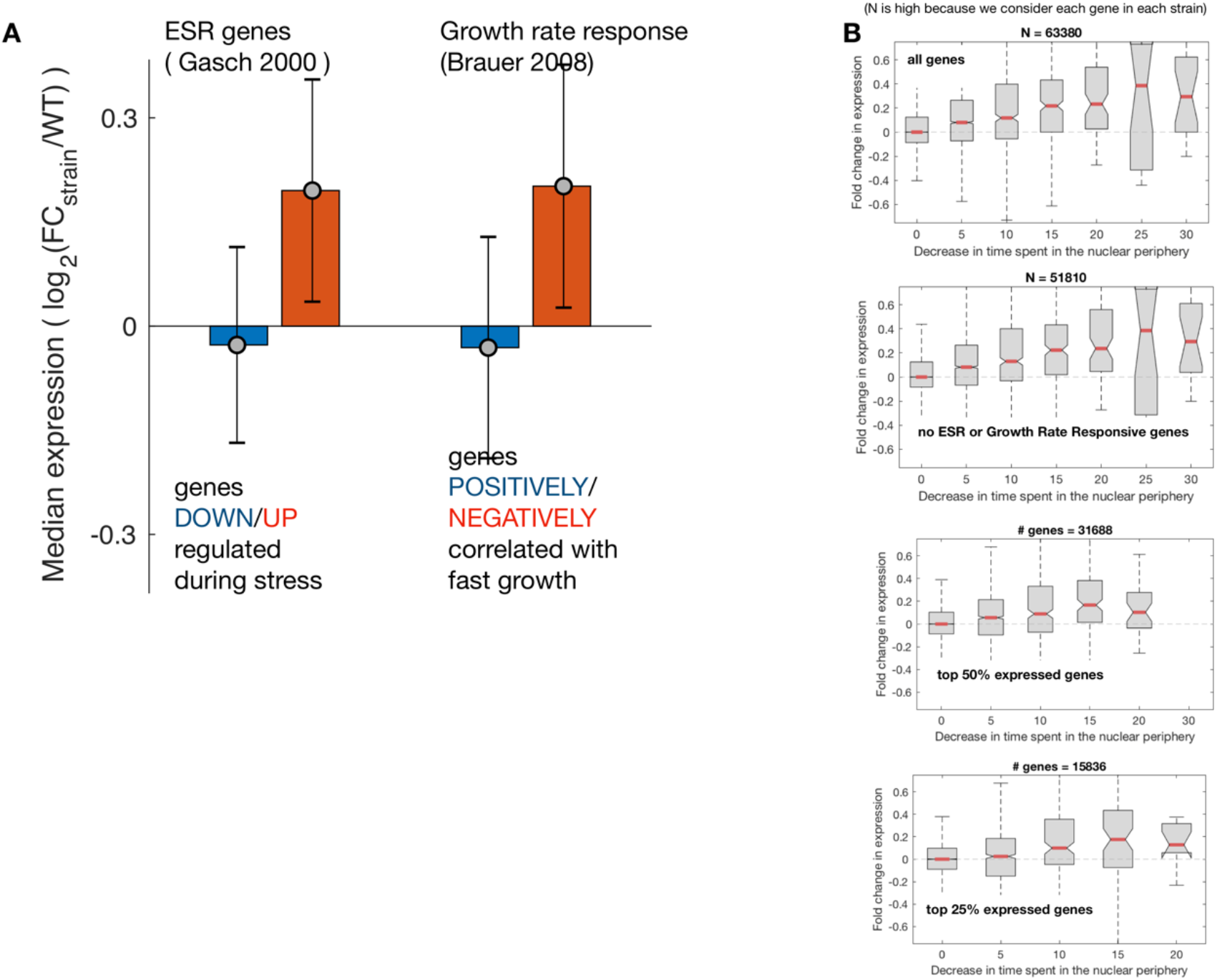
The relationship between predicted time in the nuclear periphery and change in expression remains when genes involved in the Environmental Stress Response [60] and Growth Rate Response [61] are removed. **(A)** FC strains exhibit a mild stress response. For each FC strain the genes involved in the growth rate response (right) and the ESR response (right) are differentially expressed in the FC strains, suggesting that FC strains have a minor stress phenotype. Bars show the median, error bars the standard deviation across all FC strains. **(B)** Genes predicted to spend less time in the nuclear periphery (x-axis) exhibit higher expression, even when removing ESR and growth rate responsive genes, or focusing only on highly expressed genes.

**Supplementary Figure 7.**
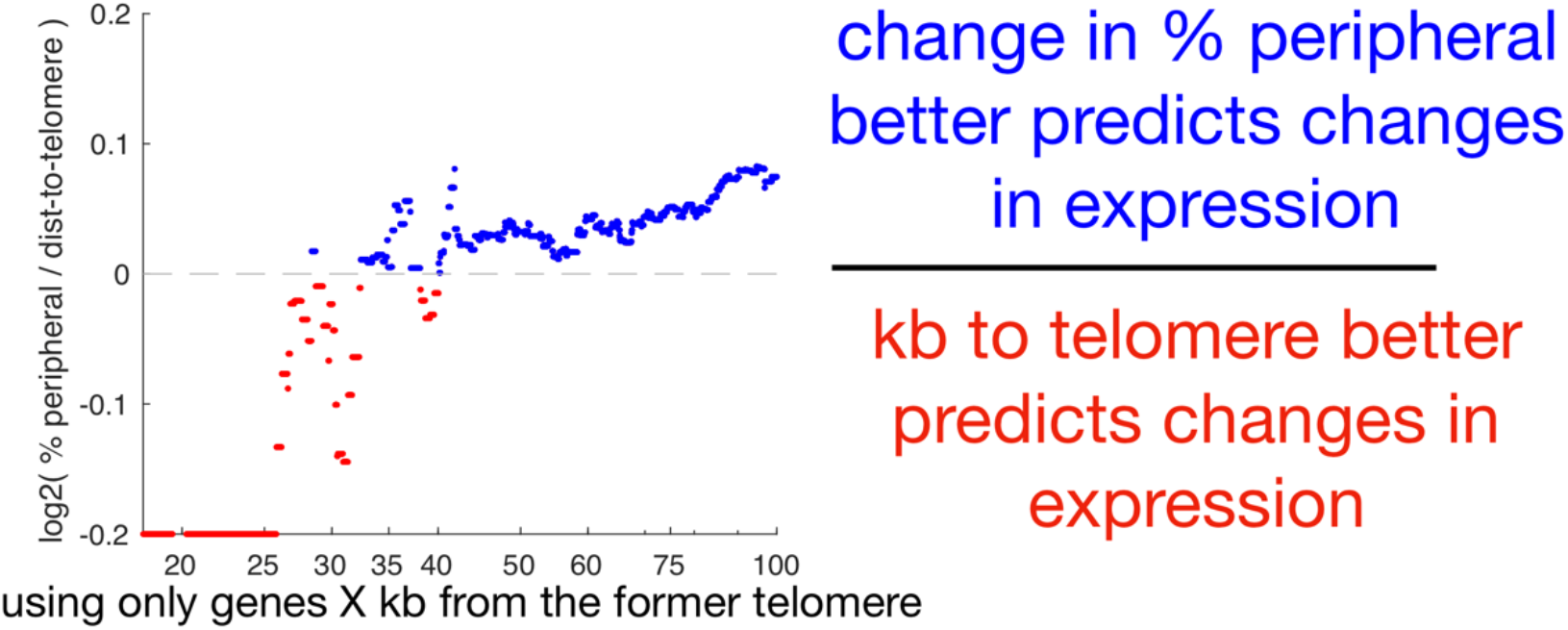
For genes further from the telomere, the predicted change in the time a gene spends in the nuclear periphery is a better predictor of changes in gene expression. Using data from all FC strains, we selected the subset of genes on chromosome arms that underwent fusion, and calculated the fold-change in expression (relative to wild-type), the change in % peripheral, and the distance to the former telomere. Each point shows the fold difference between (the ability of changes in % peripheral to predict expression) and (the ability of the log distance to the telomere to predict expression). Each value is the log2(r^2^_%peripheral_ / r^2^_dist-to-tel_) for the set of genes that are < X kb from the former telomere. Gene sets in which changes in % peripheral are better predictors of changes in expression (log2(r^2^_%peripheral_ / r^2^_dist-to-tel_) > 0) are colored blue.

**Supplementary Figure 8.**
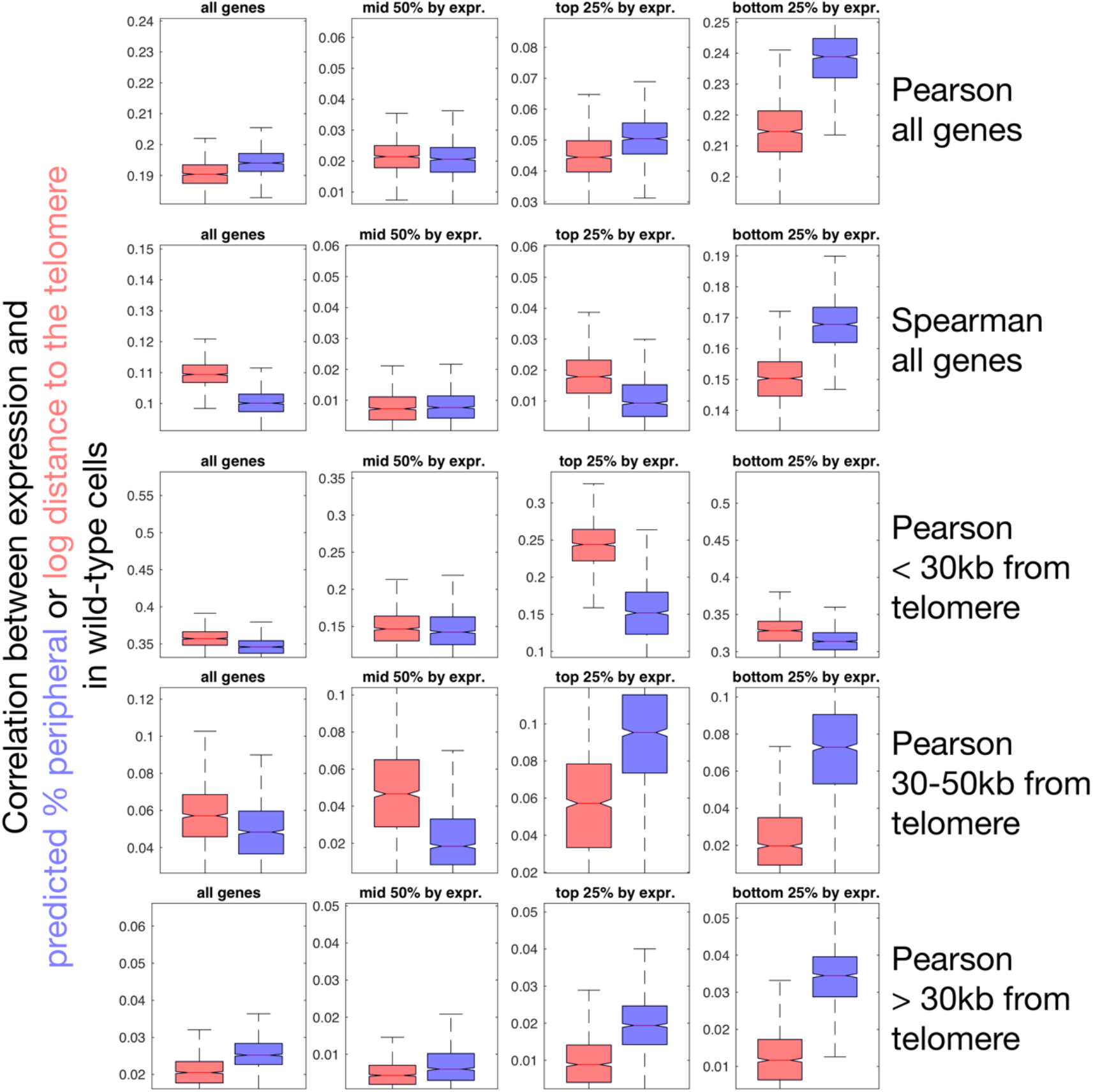
Correlation between expression and either predicted % peripheral and distance to the telomere in wild-type cells.

